# Integrative DNA copy number detection and genotyping from sequencing and array-based platforms

**DOI:** 10.1101/172700

**Authors:** Zilu Zhou, Weixin Wang, Li-San Wang, Nancy Ruonan Zhang

## Abstract

**Motivation:** Copy number variations (CNVs) are gains and losses of DNA segments and have been associated with disease. Many large-scale genetic association studies are performing CNV analysis using whole exome sequencing (WES) and whole genome sequencing (WGS). In many of these studies, previous SNP-array data are available. An integrated cross-platform analysis is expected to improve resolution and accuracy, yet there is no tool for effectively combining data from sequencing and array platforms. The detection of CNVs using sequencing data alone can also be further improved by the utilization of allele-specific reads.

**Results:** We propose a statistical framework, integrated Copy Number Variation detection algorithm (iCNV), which can be applied to multiple study designs: WES only, WGS only, SNP array only, or any combination of SNP and sequencing data. iCNV applies platform specific normalization, utilizes allele specific reads from sequencing and integrates matched NGS and SNP-array data by a Hidden Markov Model (HMM). We compare integrated two-platform CNV detection using iCNV to naive intersection or union of platforms and show that iCNV increases sensitivity and robustness. We also assess the accuracy of iCNV on WGS data only, and show that the utilization of allele-specific reads improve CNV detection accuracy compared to existing methods.

**Availability:** https://github.com/zhouzilu/iCNV

**Supplementary information:** Supplementary data are available at *Bioinformatics* online.

## 1 Introduction

Copy number variations (CNV) are large chunks of DNA that have been deleted or duplicated during evolution, leading to polymorphisms in their numbers of copies in the observed population. Studies have shown that CNV is an important type of variation in the human genome, some of which playing key roles in disease susceptibility (Freeman et al., 2006; McCarroll & Altshuler, 2007; Redon et al., 2006). Accurate identification and genotyping of CNV is important for population genetic and disease studies, and can lead to improved understanding of disease mechanisms and discovery of drug targets (Diskin et al., 2009; Glessner et al., 2009; McCarroll et al., 2008). To profile CNV, earlier studies relied on array-based technologies such as array comparative genome hybridization (CGH) or single-nucleotide polymorphism (SNP) genotyping arrays, while in recent years, next generation sequencing (NGS) technologies have allowed for high resolution CNV profiling (Abyzov, Urban, Snyder, & Gerstein, 2011; Carter, 2007; Chiang et al., 2009; Fromer et al., 2012; Jiang, Oldridge, Diskin, & Zhang, 2015; Pinkel et al., 1998; Wang et al., 2007; Zhao, Wang, Wang, Jia, & Zhao, 2013). With the drop in sequencing cost, many large-scale genetic studies have adopted whole exome sequencing (WES) and/or whole genome sequencing (WGS) to profile genetic variation in large cohorts. Often, these cohorts were previously studied using array-based technologies. For example, Alzheimer’s Disease Sequencing Project (ADSP) is an ongoing study that has 578 samples with both WGS and SNP-array data and 10913 individuals with both WES and SNP-array data; Alzheimer's Disease Genetics Consortium (ADGC) is another ongoing study that has 3084 samples with both WES and SNP-array data. It is yet uncertain in such studies how to combine data from multiple platforms, and unclear how such multi-platform integration can improve accuracy.

There is also ample room for improvement in the detection of CNV from NGS data alone. Sequencing data is subject to multiple sources of experimental noise such as GC bias and batch effects (Benjamini & Speed, 2012; Leek et al., 2010). Numerous CNV detection tools have been developed for sequencing data, but they often make contradicting detections on the same data set (Fromer et al., 2012; Klambauer et al., 2012; Krumm et al., 2012; Zhao et al., 2013). On SNP-array platforms, utilization of B-allele frequency improve CNV detection accuracy, but few of the CNV detection tools currently available for sequencing data make use of information from allele-specific reads.

Here we propose integrated Copy Number Variation caller (iCNV), a statistical framework for CNV detection that can be applied to multiple study designs: WES only, WGS only, SNP array only, or any combination of SNP and sequencing data. Compared to existing approaches, iCNV improves copy number detection accuracy in three ways: (1) utilization of B allele frequency information from sequencing data, (2) integration of sample matched SNP-array data when available, and (3) integration of improved platform-specific normalization for sequencing coverage. iCNV produces a cross-platform joint segmentation of each sample’s genome into deleted, duplicated, and normal regions, and further infers integer copy numbers in deletion and duplication regions.

To test iCNV, we first compare CNV detection accuracy using iCNV on two platforms versus simply performing intersection or union of the two platforms. Results suggest that iCNV achieves higher sensitivity and robustness. We further assess the impact of adding array data to sequencing data in CNV detection by an in silico spike-in study, and find that sensitivity increases when adding SNP array to WES, but there is negligible improvement when adding SNP array to WGS. We also consider the case where WGS is the sole platform used, and compared iCNV to other commonly used CNV calling methods on a WGS data set with pedigree information, where we find higher Mendelian concordance, indicative of higher accuracy, for iCNV detections.

## 2 Methods

### 2.1 Overview of pipeline

Figure 1 shows an overview of iCNV analysis pipeline. Input data depends on experiment design: When both SNP array and NGS data are available, the input includes (i) SNP log R ratio (LRR) and (ii) B allele frequency (BAF), which quantify, respectively, relative probe intensity and allele proportion, and (iii) sequencing mapped reads (BAM file) (Li et al., 2009). This pipeline simplifies when data from only one platform is available (Figure S1). For sequencing data, iCNV also receives target positions (BED file) for read depth background normalization. In WES, the targets are exons, while for WGS, iCNV automatically bins the genome and treats each bin as a target (the default bin size is 1kb). iCNV first performs cross-sample bias correction for sequencing data using CODEX and computes a Poisson log-likelihood ratio (PLR) for each target (Jiang et al., 2015). Heterozygous SNPs are detected and BAFs are computed within target regions using SAMTOOLS (Li et al., 2009). Integrated CNV detection is then conducted through a hidden Markov model (HMM) that treats the array intensity, array BAF, sequencing PLR and sequencing BAF as observed emissions from a hidden copy number state. The HMM segments the genome of each sample into regions of homogeneous copy number and outputs an integrated Z-score for each position that summarizes the evidence for an abnormal copy number at that position. Finally, integer-valued copy numbers are estimated in regions of high absolute Z-score, utilizing information from all platforms.

**Figure 1.**
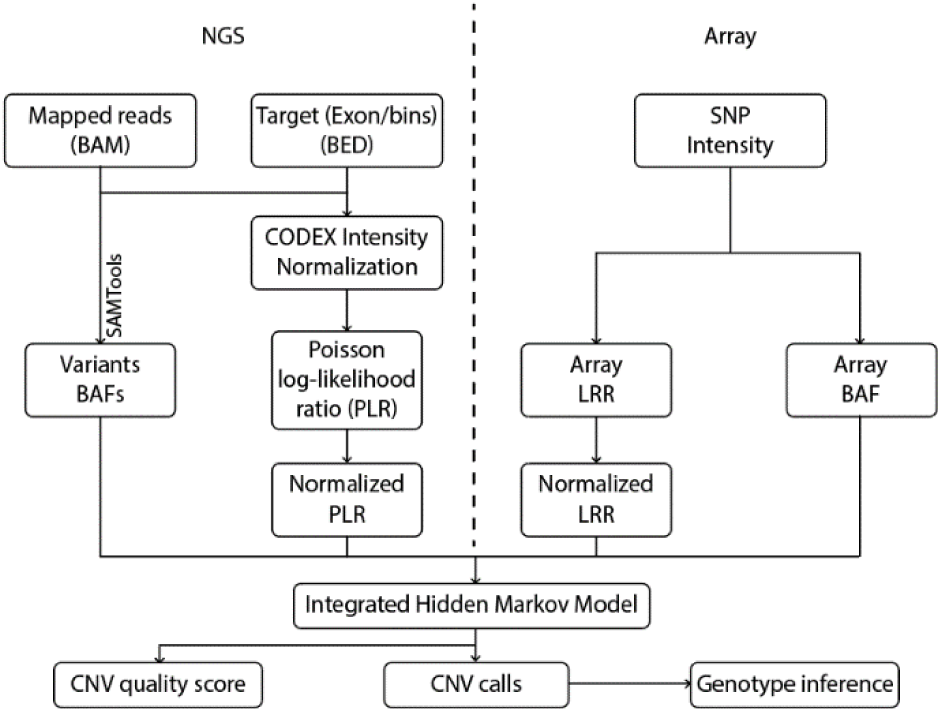
iCNV analysis pipeline including data normalization, CNV calling and genotyping using NGS and array data. For NGS data, the first step is to normalize coverage using CODEX and calculate a Poisson log-likelihood ratio (PLR), further converted to a normalized LRR by a z-transformation. The heterozygous single nucleotide positions are then found and BAF computed using SAMTools. For array data, we obtain log R ratios and BAF from raw SNP intensity data, then normalize the log R ratios. The integrated Hidden Markov Model takes these inputs and generate integrated CNV calls with quality scores. Finally, genotypes are inferred for each CNV region.

### 2.2 Platform specific normalization

Due to the heterogeneity in noise across platforms, we perform platform-specific normalization. For sequencing data, we apply CODEX (Jiang et al., 2015) normalization that removes biases related to target length, mappability, GC content and other latent systematic factors such as capture efficiency and amplification bias (Aird et al., 2011; Benjamini & Speed, 2012; Leek et al., 2010). These systematic factors are prevalent in all sequencing protocols and detrimental to CNV detection. CODEX normalization results in a PLR for each target. B allele frequencies of heterozygous variants in targets are computed by SAMTOOLs (Supplementary Method). As for array data, SNP log R ratio and BAF are standard outputs measurements, giving, respectively, the relative total probe intensity and allele proportion.

To bring SNP array intensity (*χ*_*SNP*,*j*_) and sequencing coveragederived PLR (*χ*_*WES*,*j*_ and/or *χ*_*WGS*,*j*_) to the same scale, we standardize each to produce a normalized intensity score:

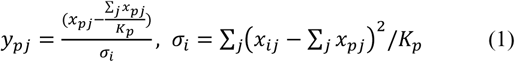

where *i* represents sample, *p* represents platform and *j* ∈ {1…*K*_*p*_} represents number of targets in the platform.

### 2.3 HMM model

After normalization, the normalized intensity score and BAFs from all platforms are analyzed by the integrated Hidden Markov Model (HMM), shown in Figure 2, which integrates evidence from all platforms and produces a joint segmentation. The HMM model has the following key features: (i) Overlapping targets (e.g. exons, bins, or SNPs) share the same underlying copy number, even if their boundaries are not identical. (ii) Each sequencing derived target (e.g. exon or genome bin) can have multiple BAFs, if multiple heterozygous SNP loci are detected in the target. In such scenarios, the BAFs are assumed independent. (iii) Exons/genome bins that don’t overlap with any heterozygous SNPs are assigned BAF value 0. (iv) For a hidden state *c*_*l*_, there are three possible values: diploid, deletion and duplication, i.e. *C*_*l*_ ∈{*del*,*dip*,*dup*}. Spe-cific integer copy numbers are inferred post-segmentation (Section 2.5.)

**Figure 2.**
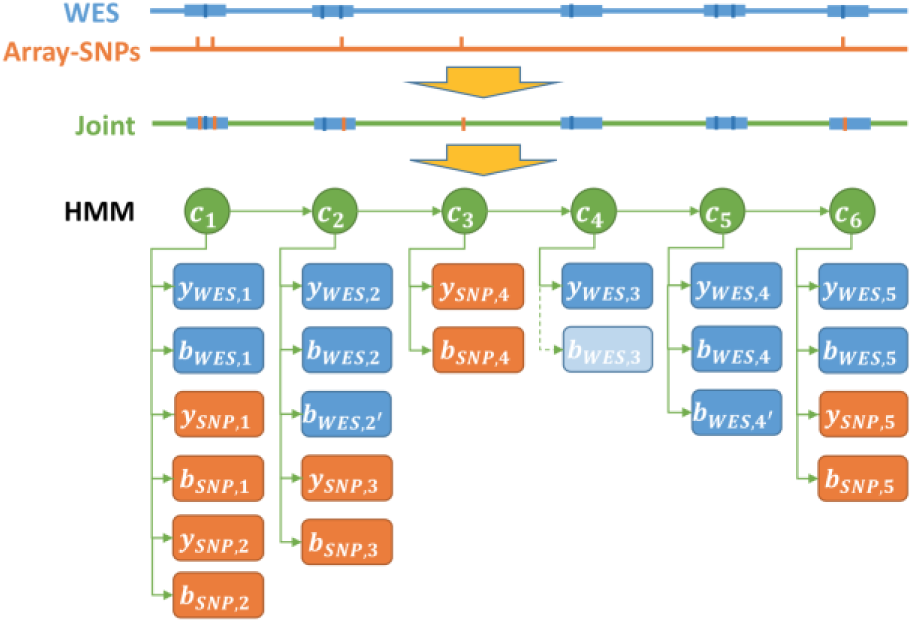
An illustration of the integrated Hidden Markov Model with array and WES data. *c*_*i*_ indicates hidden states at position *i*; *y*_*SNP*/*WES*,*i*_ indicates observation of normalized intensity score at position *i*; *b*_*SNP*_*/*_*WES*,*i*_ indicates observation of BAF at position *i*. Specifically in hidden state *c*_4_, since there is no variants BAF from WES, we assign 0.

Transition probabilities between hidden states rely on genomic distance traversed. Both XHMM (Fromer et al., 2012) and PennCNV (Wang et al., 2007) use a distance-dependent exponential attenuation factor *f*(*d*) = *e*^‒*d*/*D*^, which we also adopt in our model. *D* is set to the mean distance between targets (default 100kb). Figure S2a illustrates the relationship between distance and transition probability.

There are two types of emission distributions in the HMM, one for normalized intensity score; the other for BAF. These emission distributions are shown in Figure S2. The emission probability distribution of coverage‐ and LRR-derived normalized intensity score is a mixture of three normal centered at –*μ*_1_, 0, +*μ*_3_ that share the same standard deviation *σ*. Previously, XHMM (Fromer et al., 2012) and PennCNV (Wang et al., 2007) used a symmetric Gaussian mixture centered at –*μ*, 0, and +*μ* with standard deviation 1 representing deletion, diploid and duplication. We found that, empirically, the normalized intensity score distribution isn’t symmetric around zero and that the variance of each component varies across individuals (Figure S2c). In particular, the mean normalized intensity score for duplications is closer to 0 than that for deletions. This motivates the separate deviations, –*μ*_1_ (default value-3) for losses and +*μ*_3_ (default value 2) for gains, with shared standard deviation *σ* that is estimated by Baum-Welch separately for each sample. The emission probabilities of array‐ and sequencing-derived BAFs are more straightforward, and is a mixture of truncated normal distributions shown in Supplementary Figure S2b.

### 2.4 Parameter estimation and quality metrics

HMM parameters are fit by the Baum-Welch (EM) algorithm, which maximizes the likelihood of observed data. The Viterbi algorithm is applied to infer the most likely path given the best-fit parameters. Details are in Supplementary Method.

To quantify CNV confidence, i.e. quality score, we calculate an integrated Z-score for each hidden state as follows:

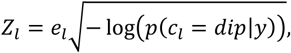

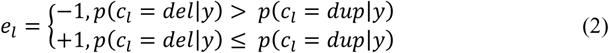

where *dip*, *dup*, and *del* are short for diploid, duplication, and deletion, respectively. The score is positive if the hidden state has higher conditional probability of being a duplication than a deletion, and negative otherwise. This quality score integrates information from intensity/coverage and BAFs across multiple platforms, allowing for easy visualization and straightforward quality assessment.

### 2.5 Copy number inference

To infer the integer copy number, we use a maximum likelihood procedure. For each CNV region, we infer deletion copy number and duplication copy number separately. Deletion can only be 0 or 1 copy. For duplication, we only characterize two cases: 3 copies or greater than 3 copies. We find it very difficult to separate copy numbers 4 or greater because (1) such events are rare, thus making it hard to infer the mean of their distributions; (2) BAF distributions among 4, 5 or more copies are too similar to be separated. The likelihood calculations are shown in the Supplementary Method. We assign the maximum likelihood copy number to each CNV region.

### 2.6 Design of spike-in

*In silico* spike-in is a useful way to assess sensitivity. Spike-in differs from simulation in that signals are inserted into real data matrices which retain the true noise structure and probe distribution, thus giving more realistic projections of detection power. In our spike-in design, we start by removing CNV regions detected by our program from original datasets, using a lenient threshold. We then add CNV signals randomly to the presumed diploid region with lengths ranging from 100bp to 500kb. Exons, bins or SNPs that overlap with the added CNV regions have their intensities and BAFs changed according to the simple and standard model described in the Supplementary Method. As a result, not all of the spike-in CNVs are detectable, especially when the data comes from WES and SNP arrays with low-resolution target set. iCNV is then applied to the spike-in dataset, using the single-platform mode for each platform as well as the integrated multi-platform mode combining the platforms. Results are compared with the underlining truth for sensitivity and precision assessment.

### 2.7 Samples and datasets

To evaluate the accuracy of iCNV, and also to serve as illustration, we analyze two sets of samples from the Alzheimer’s Disease Sequencing Project. For detailed data processing procedure see Supplementary Methods. The first set of samples comprises of 38 unrelated individuals with SNP array and WES data. The second set of samples comprises of 75 related individuals with SNP array and WGS data. The family structure in the second data set allows the assessment of detection accuracy by Mendelian concordance and the benchmarking of detection methods.

## 3 Results

### 3.1 Comparison between WES and array joint platform calls versus single platform calls

We first applied iCNV with default parameters to 38 individuals with both WES and SNP array data. We analyzed the data in three ways: Joint segmentation using both WES and array, segmentation using WES alone, and segmentation using array alone. To illustrate the relationship between the integrated HMM Z-score and the raw data input values, Figure 3 shows all of the input values along with the HMM Z-score for chromosome 22 of one typical sample. The heatmap shows detected CNV regions and integrated HMM Z-scores for this specific individual. Regions of the genome with low intensity/coverage and an enrichment of 0/1 BAFs have negative Z-scores, indicating putative deletion events; regions with high intensity/coverage and an enrichment of BAFs at duplication levels (see Supplement Method) tend to have a positive Z-score, indicating putative duplication.

**Figure 3.**
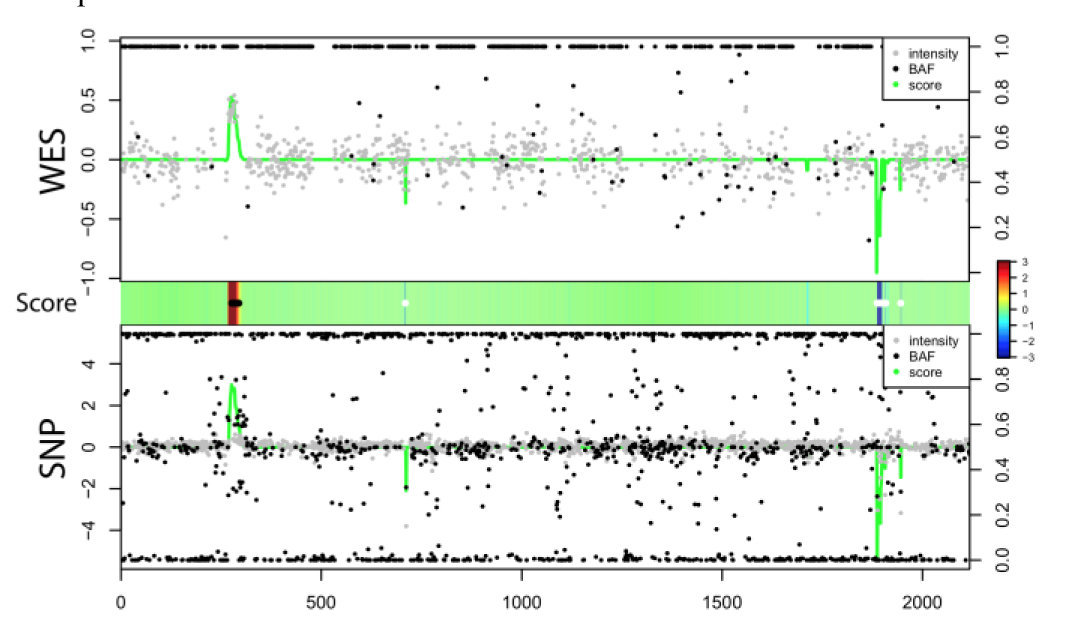
Case illustration of relationship between score (green line, range from ‐3, 3 and centered at 0, same as the middle panel), normalized intensity score (grey dots, left scale) and BAF (black dots, right scale) in WES and SNP array. Heat map in the middle indicates score and CNV calling (white dots: deletion; black dots: duplication).

Table 1 compares the integrated analysis with a simple intersection or union of results from a separate analysis of each individual platform. Details of how the union and intersections are performed are given in the Supplementary Method. In this data set, we find that an integrated analysis yields more deletion and duplications than ad hoc union of individual results from the two platforms. Compared to a simple intersection of calls from the two platforms, iCNV rescues signals that are modest in one platform while strong in another platform; an example is given in Figure S3a. Even though a simple intersection gives the most stringent call set, 87% of integrated iCNV calls have overlap with the intersection call set, implying strong confidence of the iCNV result (Table S1). On the other hand, by conventional wisdom it seems that taking a simple union of single platform calls should increase sensitivity but also increase false positive rate. However, in this case we detect more CNVs by integrated method than by simple union. Of the integrated call set, 8.94% are not present in the simple union, whereas of the union call set, 12.04% are not present in the integrated results. A signal that is moderate in both platforms would be present in the integrated call set but not in the union call set. A signal that is only present in one platform but absent in the other would be present in the union call set but not detected during integration. Compared to taking a simple union, combining the two platforms improves resolution, thus improving CNV detection power, and integration by the hidden Markov model allows one platform to “check” the calls of the other, thus improving robustness. CNVs detected by integrated analysis through iCNV, but not by simple union, are more likely to be shared across samples compared with CNVs detected by union but not integrated analysis (Figure S3b), suggesting that they are higher quality.

**Table 1.** Summary of called CNV* (WES, Array, 38 samples, Chr22)

| CNV type | WES+Array** | WES | Array | WES ∩ SNP | WES ∪ SNP |
| --- | --- | --- | --- | --- | --- |
| Deletion | 1364 | 152 | 1134 | 1252 | 1278 |
| Duplication | 381 | 124 | 144 | 252 | 268 |

\* This table measure the number of hidden state, i.e. SNP or exon/bin
\*\* Integrated CNV calling with data from both platforms

To assess the power improvement achieved by combining platforms, we conducted an in silico spike-in study. We define a detection to be a true positive if there is overlap between the detected region and a known spike-in CNV region. Power is defined as the percentage of spiked-in CNVs that are detected. We also compared detection specificity (using precision as a measure) between iCNV joint analysis and intersection or union of single platform call set. The result shows that combining SNP array with WES indeed increases power for all but the largest CNVs (Figure 5a); while not sacrificing specificity (Figure S3c).

### 3.2 Comparison between WGS and array platform calls versus individual platform calls

We next applied iCNV to 75 individuals with both WGS and array data, performing a similar analysis as above. Figure 4 shows, for a region of Chromosome 22, a heatmap of the integrated HMM Z-scores output by iCNV run on joint platform mode. Regions reported by iCNV as deletions and duplications are also shown as thick black or white lines on the heatmap. In this data set, the joint method detects more CNVs than a simple intersection of a separate analysis of the two platforms, and less than their simple union (Table 2, Figure S4a). Compared to WGS, SNP array has much lower resolution and thus detects significantly fewer CNVs. More than 71% of the SNP calls and 96% of the WGS calls overlap with iCNV joint platform calls (Table S2). On the other hand, around 92% of the joint platform calls overlap with calls made on WGS alone. Since WGS already has very good coverage across the genome, there seems to be, as expected, a much smaller power gain achievable by adding SNP arrays to the joint calling procedure.

**Table 2.** Summary of called CNV* (WGS, Array, 75 samples, Chr22)

| CNV type | WGS+Array** | WGS | Array | WGS ∩ SNP | WGS ∪ SNP |
| --- | --- | --- | --- | --- | --- |
| Deletion | 9572 | 9840 | 825 | 9235 | 10368 |
| Duplication | 5902 | 6069 | 48 | 5490 | 6090 |

\* This table measure the number of hidden state, i.e. SNP or exon/bin
\*\* Integrated CNV calling with data from both platforms

**Figure 4.**
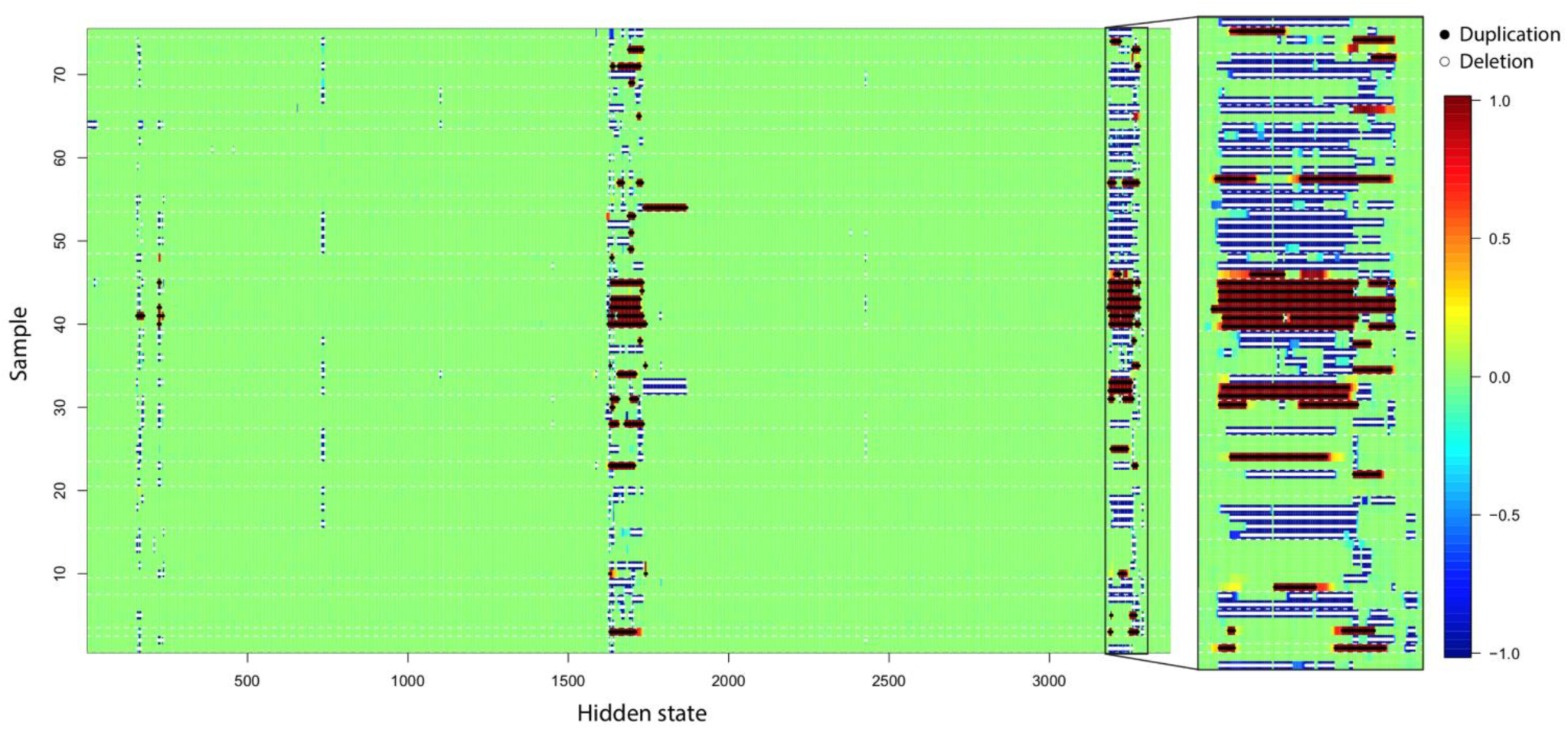
CNV score at each hidden state from 75 related individuals with SNP array and WGS data on Chr22 CNV hot region (17000-20800kb). Region (20320-20720kb) is enlarged for detailed visualization.

We conduct another in silico spike-in study to study the power gains from adding array data to WGS. The result shows that WGS has comparable power to the integrated method, performing only slightly worse (Figure 5b). However, for the detection of small CNVs in SNP-dense regions, SNP-array does add valuable information. Precision, reflecting specificity, of joint detection is similar to the intersection method, and sensitivity is similar to the union method (Figure S4c). Thus, iCNV achieves the best of both worlds: The high precision of a stringent intersection procedure as well as the high sensitivity of a relaxed union procedure.

**Figure 5.**
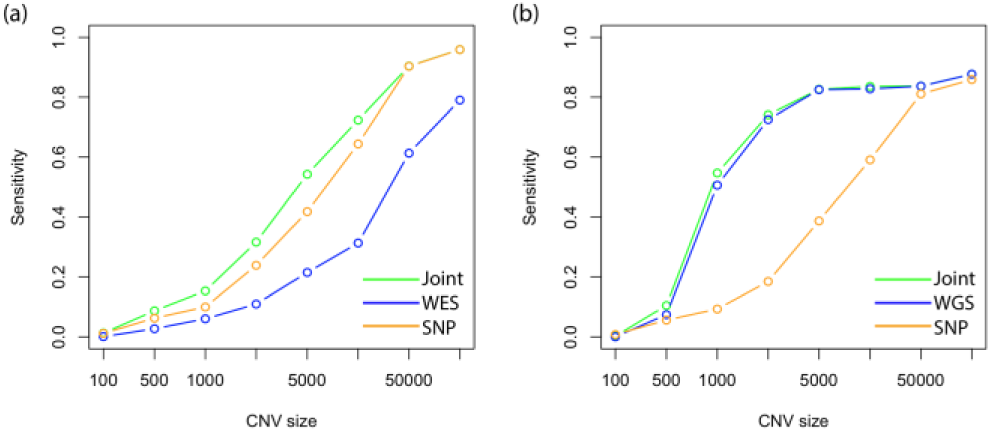
Relationship between power (or sensitivity) and size of CNV using spike-in in (a) WES and SNP array scheme and (b) WGS and SNP array scheme. Joint means integrated analysis using iCNV with both WES and SNP array data.

### 3.3 Improvement of CNV detection by WGS alone

We utilized pedigree data that is available for the 75 individuals with WGS to assess the improvement of iCNV over existing methods for CNV detection on WGS alone. True germline CNVs are more likely to be shared between related individuals than between unrelated individuals, whereas false positives are not expected to have enriched sharing between related. Based on this fact, we can compute the CNV sharing frequency *f* at each position *l* between related individuals, defined as

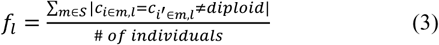

where S represents the set of families, against the cohort call frequency, defined simply as the fraction of individuals where this CNV is detected among the 75 individuals analyzed. As a baseline for comparison, we compute the expected sharing frequency under random permutation of family labels, which represents the null scenario of a random detection. Enrichment of detections above the permutation-derived mean is evidence for enrichment of true positives.

Since these samples also have array data, a comparison of calls made by integrating WGS and SNP array versus calls made by a separate analysis of each platform alone is shown in the Supplementary Method. Based on this metric, there is no detectable gain of adding SNP array to WGS, which is expected given our analysis in Section 3.2 showing very little power gain from adding SNP array to WGS. In addition, CNVs detected by integrated analysis but not union are slightly more likely to share within family compared with CNVs detected by union but not integrated analysis (Figure S4b), indicating the iCNV calls are more accurate than a simple union.

Using this pedigree-based quality metric, we compared iCNV, run on WGS-only mode, to the WGS-based CNV detection methods CNVnator (Abyzov et al., 2011) and cn.MOPS (Klambauer et al., 2012). CNVnator and cn.MOPS were run with default parameters and the same bin size (1kb) as iCNV. Results are shown in Figure 6 and Table 3. To quantify the enrichment of familial sharing, we computed a sharing enrichment score by subtracting from the observed familial sharing frequency its permutation mean and then dividing the difference by its permutation standard deviation; see Supplementary Method for details. Larger sharing enrichment score indicates stronger within family enrichment while zero indicates no enrichment. As shown in Figure 6, cn.MOPS and iCNV show significant enrichment of familial sharing as compared to CNVnator. The enrichment scores of the iCNV call set are significantly higher than CNVnator (one-side t.test, pvalue= 1.28 × 10^‒7^) and cn.MOPS (one-side t.test, pvalue=1.54 × 10^‒2^).

**Figure 6.**
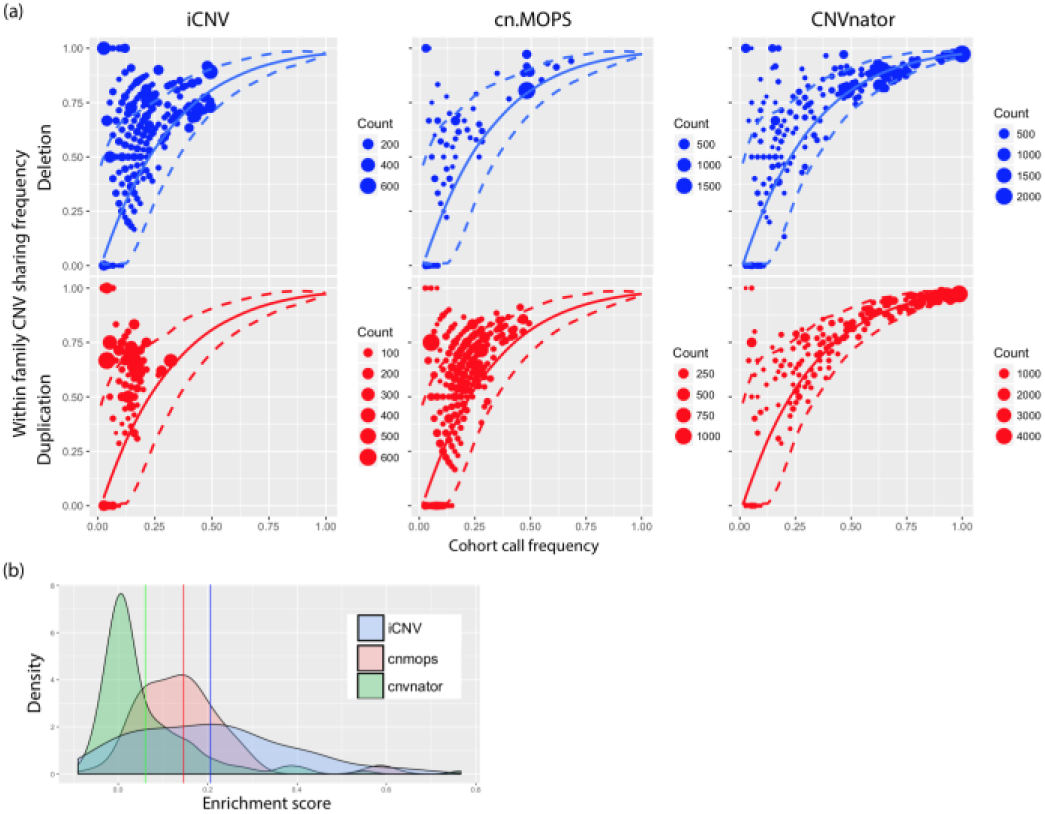
(a) CNV detection assessment of iCNV, cn.MOPS, CNVnator by real dataset with pedigree information in WGS only scheme. Plot shows the enrichment of CNVs (deletion or duplication) with in a pedigree. The solid line and the dash lines are the null distribution mean and 95% confidence interval, respectively, calculated by permutation. Size of the dots shows number of CNV events counts. (b) Enrichment score distribution plot between iCNV, cn.MOPS and CNVnator. Larger number indicates higher enrichment. Solid line is the distribution mean for each method.

**Table 3.**
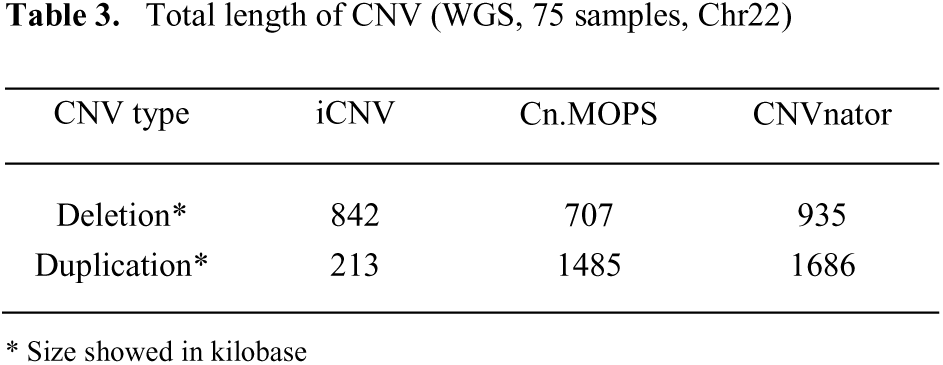
Total length of CNV (WGS, 75 samples, Chr22)

| CNV type | iCNV | Cn.MOPS | CNVnator |
| --- | --- | --- | --- |
| Deletion* | 842 | 707 | 935 |
| Duplication* | 213 | 1485 | 1686 |

\* Size showed in kilobase

### 3.4 CNV genotype inference

An integer copy number is assigned to each CNV region after the detection step. The assignment is based on maximizing a likelihood function that quantifies the probability of the observed normalized intensities and BAFs for each copy number state. The data input (normalized intensities and BAFs from all platforms) are shown in Figure 7, along with their copy number assignments shown as contours. The marginal densities of the normalized intensity values for each platform seem to be well modeled by a mixture of normals with platform-specific mean and variance. The BAF is much noisier and do not show any platform specific trend. Thus, the likelihood model we use is based on a mixture of normal for the normalized intensities, with platform and copy number specific means determined by an initial K-means clustering step, and a mixture of truncated normal for the BAF with pre-fixed means and standard deviations. The maximum likelihood copy number state is assigned to each segment.

**Figure 7.**
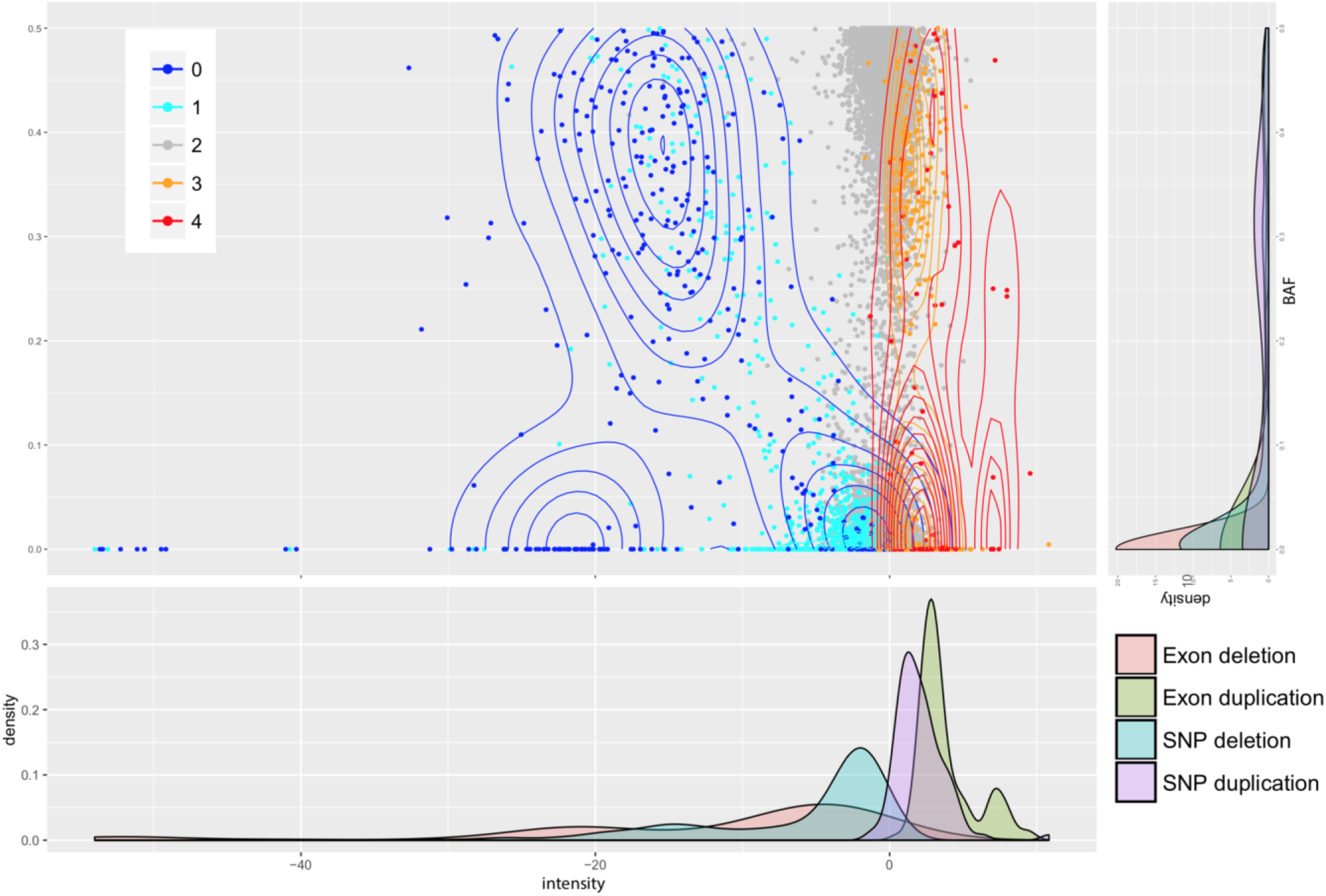
Raw intensity score and BAFs distribution of CNV genotype inference. Contour plot shows distribution density. Each margin respectively shows density distribution of intensity score and BAF in each CNV genotype category.

## 4 Discussion

We have proposed a method, iCNV, to improve CNV detection and genotyping accuracy using high throughput sequencing data, allowing for integration of SNP-array data. The distinguishing features of iCNV compared with existing methods are as follows: *(i)*iCNV adopts CODEX to improve the normalization of sequencing data, removing biases due to target length, mappability, GC content and other latent systematic factors; *(ii)* iCNV utilizes B-allele frequency information from sequencing data, which is valuable for CNV detection and exact copy number inference; *(iii)* Array data, if available, are combined with sequencing data to allow more sensitive and robust CNV detection than either platform alone; *(v)* iCNV outputs a Z-score from an integrated HMM that summarizes evidence across multiple platforms, allowing for easy visualization and quality assessment; *(vi)* Even though we combine cross-platform data for CNV detection, we use platform-specific parameters for exact copy number estimation and thus minimize noise effect due to platform specific latent variables.

How much does SNP-array data add to NGS data for CNV detection? Our results, based on spike-ins and pedigree-based quality evaluations, show that SNP-arrays give a significant boost in accuracy to WES but relatively little gains for WGS. For SNP-array data, although we applied Illumina platform as example, we could incorporate any data as long as both Log R ratio and B allele frequency are available. For CNV detection and genotyping using WGS alone, we compared iCNV against other read depth based CNV detection methods including cn.MOPS (Klambauer et al., 2012) and CNVnator (Abyzov et al., 2011). Germline CNVs detected by iCNV have higher within-family sharing than the other methods being compared, suggesting a higher accuracy.

For WGS data, bin length affects analysis results. In the trade-off between sensitivity and specificity, larger bin size increases specificity while decreasing sensitivity. We have found 1kb to be a good default value. Users can customize longer or shorter bins depend on analysis goal and sequencing coverage. iCNV is implemented in R and is available on Bioconductor (under revision) and Github (https://github.com/zhouzilu/iCNVhttps://github.com/zhouzilu/iCNV). For chromosome 22 on 75 samples, the run time for joint detection (WGS and SNP-array) is less than 5 minutes on a 16GB-RAM laptop. Runtime scales linearly with number of samples and genome size. iCNV allows parallelization across samples once the CODEX normalization step is finished, and the entire procedure can be parallelized across chromosomes. Collectively, iCNV provides a systematic framework and an efficient, scalable toolset for single and cross platform CNV detection.

## Acknowledgements

Subjects recruitment and sample/phenotype collection were conducted by Alzheimer's Disease Centers (ADC), National Cell Repository for Alzheimer's Disease (NCRAD) and The National Alzheimer's Coordinating Center (NACC). The Alzheimer's Disease Genetics Consortium (ADGC) generated SNP array genotype data. The Alzheimer’s Disease Sequencing Project (ADSP) produced sequencing data. We thank contributors who collected samples used in this study, as well as patients and their families, whose help and participation made this work possible.

